# Specialized coding patterns among dorsomedial prefrontal neuronal ensembles predict conditioned reward seeking

**DOI:** 10.1101/2020.12.14.422672

**Authors:** Roger I. Grant, Elizabeth M. Doncheck, Kelsey M. Vollmer, Kion T. Winston, Elizaveta V. Romanova, Preston N. Siegler, Christopher W. Bowen, James M. Otis

**Author notes:** Correspondence should be addressed to Dr. James M. Otis at.

## Abstract

Non-overlapping cell populations within dorsomedial prefrontal cortex (dmPFC), defined by gene expression or projection target, control dissociable aspects of reward seeking through unique activity patterns. However, even within these defined cell populations considerable cell-to-cell variability is found, suggesting that greater resolution is needed to understand information processing in dmPFC. Here we use two-photon calcium imaging in awake, behaving mice to monitor the activity of dmPFC excitatory neurons throughout Pavlovian sucrose conditioning. We characterize five unique neuronal ensembles that each encode specialized information related to a reward, reward-predictive cues, and behavioral responses to reward-predictive cues. The ensembles differentially emerge across learning – and stabilize after learning – in a manner that improves the predictive validity of dmPFC activity dynamics for deciphering variables related to behavioral conditioning. Our results characterize the complex dmPFC neuronal ensemble dynamics that relay learning-dependent signals for prediction of reward availability and initiation of conditioned reward seeking.

## INTRODUCTION

The dorsomedial prefrontal cortex (dmPFC) has garnered considerable interest due to its dysregulation in diseases associated with disordered reward processing (Chen et al., 2018; Courchesne et al., 2011; Dienel and Lewis, 2019; Holmes et al., 2018; Koob and Volkow, 2010; Ye et al., 2012). These abnormalities include aberrant cell morphology and regional mass (Courchesne et al., 2011), abnormal activity patterns (Dienel and Lewis, 2019), and reduced behavioral performance on tasks that involve dmPFC activity (Goldstein and Volkow, 2011). Despite this knowledge, how unique cell types in dmPFC encode complex reward-related information to guide behavioral output is unclear, limiting our understanding of how reward processing occurs in healthy individuals as compared to those with neuropsychiatric diseases.

Neuronal activity in dmPFC neurons has been observed to be heterogeneous in a variety of behavioral tasks (Kim et al., 2016; Kobayashi et al., 2006; Matsumoto et al., 2003; Powell and Redish, 2014), including those that involve reward-seeking behaviors (Sun et al., 2011; Moorman and Aston-Jones, 2015; Otis et al., 2017, 2019; Sparta et al., 2014). Recent studies have aimed to resolve this heterogeneity through cell-type specific recording strategies, such as *in vivo* calcium imaging in genetically or projection-defined neurons (Otis et al., 2017, 2019; Siciliano et al., 2019; Ye et al., 2016). Although some variability can be explained by identified neuronal subpopulations, a vast majority of response diversity remains unexplained (Otis et al., 2017). For example, we recently demonstrated that many dmPFC excitatory neurons that project to the nucleus accumbens (NAc) show learning-related responses to reward-predictive cues, with two-thirds of the responding cells being excitatory responders and the other third being inhibitory responders. Similarly, dmPFC neurons that project to the paraventricular thalamus (PVT) also show learning-related responses to reward-predictive cues, although about two-thirds of responding cells are inhibitory responders and the other one-third are excitatory responders (Otis et al., 2017, 2019). Finally, responses can be further subdivided by their temporal relation to the cue, as well as the reward (i.e., anticipation versus consumption). Overall, although subpopulations of dmPFC output neurons could be labeled as ‘generally excited’ (e.g., dmPFC NAc) or ‘generally inhibited’ (e.g., dmPFC PVT), such assignment ignores much of the variability that is likely critical for behavioral control. Thus, a more thorough and unbiased means of defining the heterogeneous activity patterns among dmPFC output neurons is needed to understand how these neurons are engaged during reward-related behavioral tasks.

Here we use *in vivo* two-photon calcium imaging to measure and longitudinally track the activity dynamics of single dmPFC excitatory output neurons throughout a Pavlovian sucrose conditioning task. We observe five unique neuronal ensembles that encode specialized information related to the sucrose reward, sucrose-predictive cues, and behavioral responses to those cues. These five ensembles differentially emerge across learning in a manner that improves the predictive validity of dmPFC population dynamics for deciphering reward delivery, cue presentation, and behavior. Finally, we find that single neurons within dmPFC neuronal ensembles display day-to-day stability after learning. Overall, we find that heterogeneous excitatory neuronal ensembles in dmPFC evolve specialized coding patterns across cue-reward learning that are maintained after learning. Our results highlight the importance of ensemble-specific recording and manipulation strategies for understanding the function of dmPFC activity for reward processing.

## METHODS

### Subjects

Male and female C57BL/6J mice (8 wks old/20 g minimum at study onset; Jackson Labs) were group-housed pre-operatively and single-housed post-operatively under a reversed 12:12-hour light cycle (lights off at 8:00am) with access to standard chow and water *ad libitum*. Experiments were performed in the dark phase and in accordance with the NIH Guide for the Care and Use of Laboratory Animals with approval from the Institutional Animal Care and Use Committee at the Medical University of South Carolina.

### Surgeries

Mice were anesthetized with isoflurane (0.8-1.5% in oxygen; 1L/minute) and placed within a stereotactic frame (Kopf Instruments) for cranial surgeries. Ophthalmic ointment (Akorn), topical anesthetic (2% Lidocaine; Akorn), analgesic (Ketorolac, 2 mg/kg, ip), and subcutaneous sterile saline (0.9% NaCl in water) treatments were given pre- and intra-operatively for health and pain management. Before lens implantation, a virus encoding the calcium indicator GCaMP6s (AAVdj-CaMK2α-GCaMP6s; UNC Vector Core) was unilaterally microinjected into the dmPFC (specifically targeting prelimbic cortex; 400nl; AP, +1.85mm; ML, −0.50mm; DV, −2.45mm). Next, a microendoscopic GRIN lens (gradient refractive index lens; 4mm long, 1mm diameter; Inscopix) was implanted dorsal to dmPFC (AP, +1.85mm; ML,-0.50mm; DV,-2.15mm) as previously described (Otis et al., 2017; Resendez et al., 2016). A custom-made ring (stainless steel; 5 mm ID, 11 mm OD) was then adhered to the skull using dental cement and skull screws. Head rings were scored on the base using a drill for improved adherence. Following surgeries, mice received antibiotics (Cefazolin, 200 mg/kg, sc), and were allowed to recover with access to food and water *ad libitum* for at least 21 days. Histology was performed after the experiments to ensure virus placement in dmPFC and lens placement dorsal to dmPFC GCaMP6s-expressing neurons.

### Behavioral procedure

Following recovery from surgery, mice were water restricted (water bottles removed from cages), and 0.6-2.0 mL of water was delivered every day to a dish placed within each home cage. Mice were weighed daily and given the appropriate amount of water to maintain ~90% of their initial body weight. No health issues related to dehydration arose at any point during or after implementation of this protocol. Once mice reached ~90% of their free drinking weight, they underwent 3 days of 30-minute habituation sessions, during which they were head-restrained and received droplets of sucrose (12.5% sucrose in water; ~2.0 μl) at random intervals through a gravity-driven, solenoid-controlled lick spout (see Figure 1A). Next, mice underwent head-fixed Pavlovian conditioning, wherein two conditioned stimuli (CS; 70 dB; 3 kHz or 12kHz as described in Otis et al., 2017) were randomly presented 50 times each (see Figure 1B). One tone (CS+) was paired to the delivery of a sucrose reward after a one second trace interval, whereas the other tone (CS-) did not result in sucrose delivery. The trace interval was included to allow isolated detection of sensory cue- and sucrose reward-related neuronal activity patterns, as described previously (Otis et al., 2017). The inter-trial interval between the previous reward delivery (CS+) or omission (CS-) and the next cue was chosen as a random sample from a uniform distribution ranging from 20 seconds to 50 seconds. Cue discrimination was quantified using the area under a receiver operating characteristic (auROC) formed by the number of baseline-subtracted licks during the CS+ versus CS- trace intervals. For all behavioral experiments, we classified sessions as ‘early’ or ‘late’ in learning based on animals’ behavioral performance (early, any sessions before auROC > 0.65; late, any sessions after auROC > 0.66).

**Figure 1.**
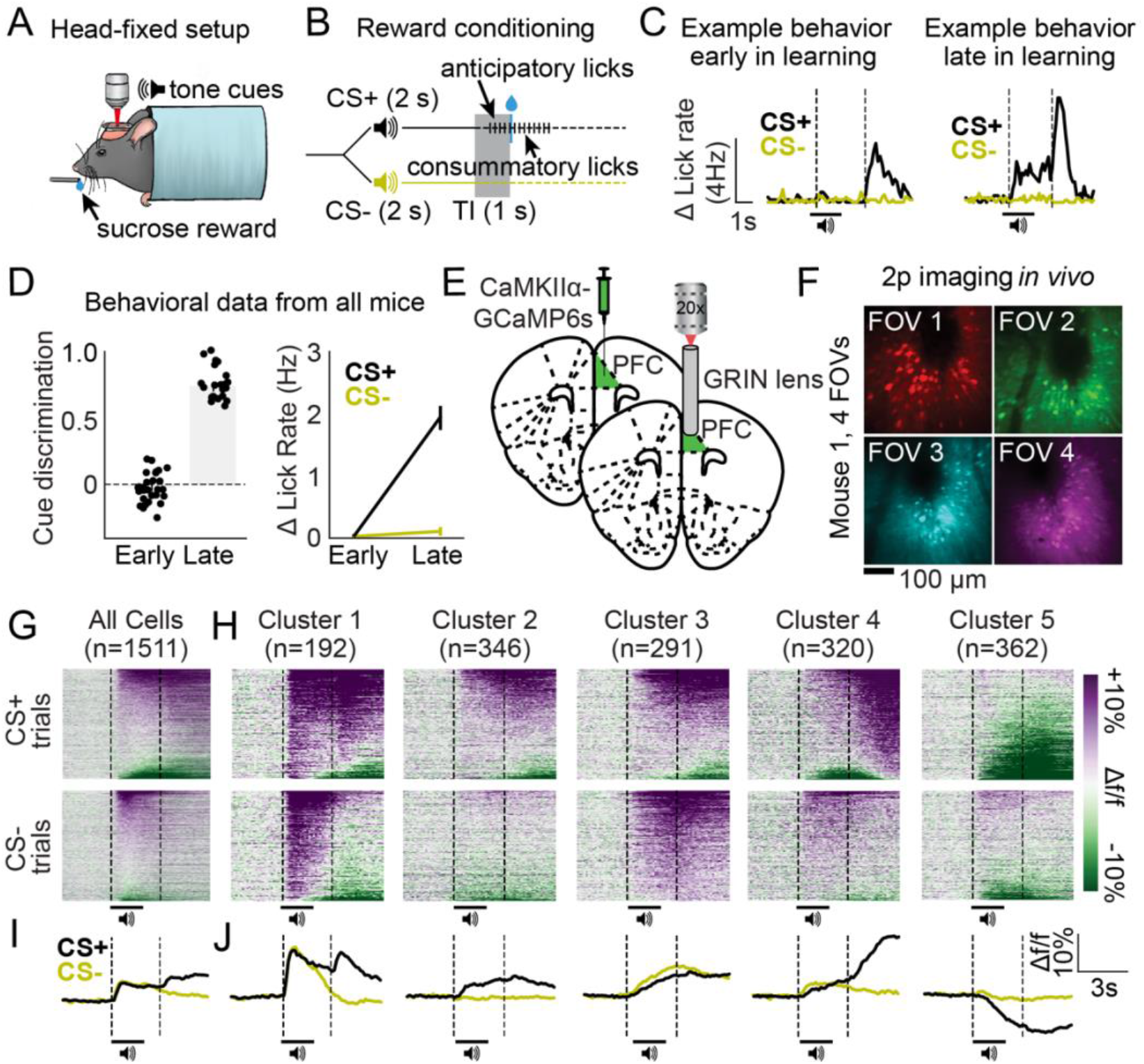
Distinct excitatory neuronal ensembles revealed in dmPFC during head-fixed Pavlovian conditioning. (**A**) Illustration of head-fixation for sucrose conditioning experiments with concurrent 2p imaging. (**B**) Schematic for sucrose conditioning experiments, in which the CS+ and CS-are presented in a random order 50 times each. The CS+ denotes the availability of a liquid sucrose reward following a 1s trace interval (TI). Anticipatory licks are seen during the trace interval in well-trained mice. (**C**) Example behavior data early in learning (left) versus late in learning (right). (**D**) Cue discrimination scores (auROC; CS+ vs. CS-) and change in lick rate for each cue during early and late in learning behavioral sessions. (**E-F**) Surgery schematic (E) allowing for *in vivo* imaging of GCaMP6s-expressing neurons (F). (**G-H**) Heat maps displaying the responses of all dmPFC neurons (G) and responses separated by cluster (H) aligned to the cues. (**I-J**) Line plots displaying the mean activity traces of all cells (I) and mean activity of all cells separated by cluster (J).

### Multiphoton imaging

We visualized and longitudinally tracked GCaMP6s-expressing dmPFC neurons throughout Pavlovian sucrose conditioning using a multiphoton microscope (Bruker Nano Inc.) equipped with: a hybrid scanning core with galvanometers and fast resonant scanners (>30 Hz; we recorded with 4 frame averaging to improve spatial resolution), GaAsP photodetectors with adjustable voltage and gain, a single green/red NDD filter cube, a long working distance 20x air objective designed for optical transmission at infrared wavelengths (Olympus, LCPLN20XIR, 0.45NA, 8.3mm WD), a moveable objective in the X, Y, and Z dimensions, and a tunable InSight DeepSee laser (Spectra Physics, laser set to 920nm, ~100fs pulse width). Data were acquired using PrairieView software and converted into an hdf5 format for motion correction using SIMA (Kaifosh et al., 2014). Fields of view (FOVs) were selected prior to “early” behavioral imaging sessions, and each FOV was separated by at least 50μm in the Z-plane to avoid visualization of the same cells in multiple FOVs. Within each FOV, regions of interest around each cell were manually traced using the ‘polygon selection’ tool in FIJI (Schindelin et al., 2012). Care was taken to only assign regions of interest to visually distinct cells. In cases where neighboring cells or processes overlapped, regions of interest were drawn to exclude areas of overlap. Fluorescence trace extraction and all subsequent analyses were performed using custom-written Python code (Namboodiri et al., 2019; Otis et al., 2019).

### Data collection and statistics

Behavioral sessions were controlled through a custom MATLAB graphical user interface connected to Arduino and associated electronics. Transistor-transistor logic (TTL) pulses between the Arduino and the microscope were used to start and stop imaging and behavioral programs, and to allow frame timestamp collection for post-hoc synchronization of the behavioral and imaging data. Behavioral data were recorded and extracted using MATLAB, analyzed and graphed using Python, and figures were produced using Adobe Illustrator. Behavioral data were presented as normalized auROC “cue discrimination” scores (2 x (auROC-0.5)), comparing licking rates during the one second period between each CS+ and reward (1s trace interval) or CS- and reward omission (1s). A cue discrimination score of −1.0 would therefore suggest more licking during all CS-trials versus CS+ trials. In contrast, a score of +1 would suggest more licking on all CS+ trials versus CS-trials. Following data collection, independent t-tests were used to compare behavioral data during all imaging/behavioral sessions across days (e.g., early in learning vs. late in learning). Additionally, a two-way ANOVA was used to compare baseline-subtracted lick rates (Δ lick rate; calculated as: 1 second trace interval licking frequency – 3 second baseline licking frequency), followed by Bonferroni multiple comparisons tests if appropriate (Otis et al., 2017).

Fluorescence signals from each cell were extracted from the motion-corrected data using custom-written Python code. Activity in each cell was then aligned to the epoch containing a ~3-second baseline (23 frames), ~3-second cue period (including CS+ or CS- trace intervals; 23 frames), and ~3-second reward period (following sucrose delivery or omission; 23 frames). This resulted in 69 frames for each CS+ and CS- trial, which was then combined into a 138-column vector of data points for each cell, referred to as a peristimulus time histogram (PSTH). Due to the robust responses of dmPFC neurons late in learning (as also seen in Otis et al., 2017), data included within a 138 × 1511 vector (138 frames, 1511 neurons) from late in learning sessions then underwent principal components analysis to reduce their dimensionality in preparation for clustering, an unbiased means of identifying putative neuronal ensembles in dmPFC. We used an analysis and code that was previously created by others and kindly shared (full description of clustering in Namboodiri et al., 2019). The principal components were determined using the point of inflection on a scree plot, which plots the PSTH variance explained versus an increasing number of principal components. Beyond this inflection point, minimal variability can be explained by additional principal components. The data were then projected onto the subspace formed by these principal components, which was fed into the clustering algorithm. We used the Scikit-learn function *sklearn.cluster.SpectralClustering* to perform spectral clustering on these data, which uses a k-nearest neighbor connectivity matrix to create clusters. The optimal number of clusters and nearest neighbors were determined by checking a range of values for each and choosing the parameters with the maximum silhouette score. Silhouette scores provide a measure of how well the data “fit” into each cluster. After clustering, each neuron was assigned a label based on its corresponding cluster.

We compared the behavioral performance of each mouse with the number of neurons detected per ensemble in the corresponding FOV visualized during that behavioral session. Specifically, we used Pearson correlations to compare cue discrimination scores (auROC, CS+ vs. CS-) with the number of neurons detected in each ensemble during late in learning behavioral sessions (one unique FOV per session; Figure 2A). Additionally, we used Pearson correlations to compare the initiation of licking “errors”, wherein mice increased licking rates following the presentation of the CS-(auROC, CS-vs. baseline epoch), with the number of detected neurons per ensemble (Figure 2B).

**Figure 2.**
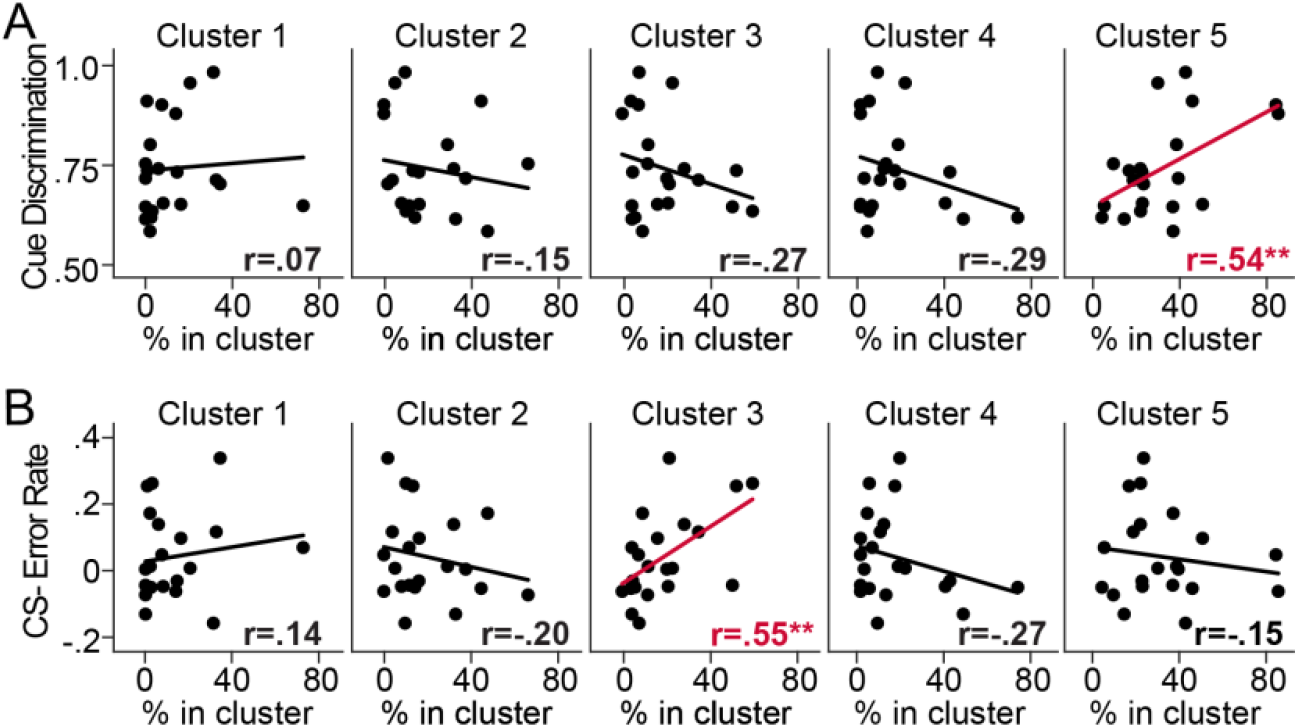
Behavioral performance is predicted by the relative percentage of dmPFC excitatory neurons within select ensembles. (**A**) Correlation plots separated by cluster displaying the relationship between cue discrimination behavioral scores after learning (auROCs: CS+ vs. CS-licking) and the percentage of detected neurons in a cluster during that behavioral session. The relative percentage of neurons in Cluster 5 positively predicted behavioral performance (\*\**p* = 0.01). (**B**) Correlation plots separated by cluster displaying the relationship between CS-licking error rate after learning (auROC: CS-vs. baseline licking) and the percentage of neurons in a cluster during that behavioral session. The percentage of neurons in Cluster 3 positively predicted CS-licking error rate (\*\**p* < 0.01).

Decoding analyses were employed to determine whether neuronal activity within each FOV, and each ensemble within each FOV, could predict variables within the behavioral paradigm better than chance. Specifically, we used these neuronal data to inform a decoder to predict (1) <u>CS+</u>: 2s CS+ epoch vs. 2s baseline, (2) <u>CS-</u>: 2s CS-epoch vs. 2s baseline, (3) <u>CS+ vs. CS-</u>: 2s CS+ epoch vs. 2s CS-epoch, (4) <u>Reward</u>: 1s epoch starting at sucrose delivery vs. 1s pre-sucrose baseline, and (5) <u>Licking</u>: relative licking rate for each mouse during CS+ trials; 6s epoch starting from CS+ onset to include anticipatory licking and sucrose consumption. Decoding was performed on the entire dataset (Population Decoding, Figure 3A-B), and separately for each ensemble (Ensemble Contribution, Figure 3C-D). To perform these analyses, we used a binary decoder as described previously (Otis et al., 2017, 2019), implemented using the Scikit-learn functions *sklearn.discriminant_analysis*, *sklearn.svm*, and *sklearn.decomposition*. For population decoding (decoding based on the activity of all neurons within each FOV), decoding scores were normalized to the maximum value observed for a decoded variable (which was CS+ vs CS-, late in learning). For ensemble-specific decoding (decoding based on the activity of all neurons within each ensemble within each FOV), scores were normalized to the maximum for each particular variable, such that the contribution of each ensemble to variable decoding could be evaluated. To determine whether decoding performance was significantly better than chance, we compared the decoding accuracy to that of randomized “shuffled” data using two-way ANOVAs (population decoding) or one-way ANOVAs (ensemble contribution), followed by program-recommended post-hoc analysis (Dunnett’s for one-way ANOVA, Tukey’s for two-way ANOVA). Because shuffled data were not significantly different between ensembles, these data were combined to improve data visualization and simplify analysis (dotted lines in Figure 3C).

**Figure 3.**
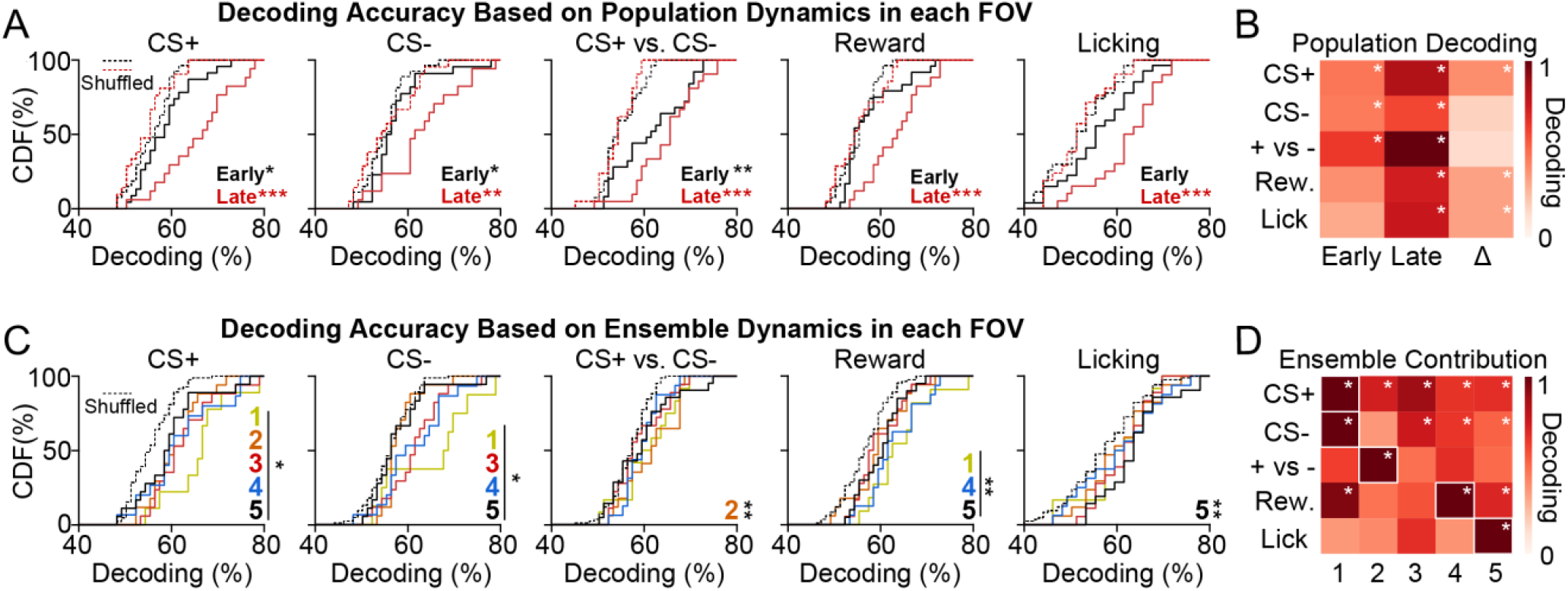
Activity of dmPFC excitatory neuronal ensembles can decode specialized information related to reward seeking. (**A**) CDF plots illustrating the population decoding accuracy for variables related to conditioned reward seeking (CS+, CS-, CS+ vs. CS-, reward, and licking). Dotted lines refer to shuffled control data for early and late in learning. (**B**) Heat maps depicting population decoding accuracy early in learning (1^st^ column), late in learning (2^nd^ column), and the change across learning (3^rd^ column). Data have been normalized to CS+ vs. CS- late in learning to provide comparison of decoding strength across variables. (**C**) CDF plots illustrating the decoding accuracy of specific ensembles for variables related to conditioned reward seeking. Dotted lines refer to shuffled control data for all ensembles. (**D**) Heat maps depicting the contribution of each ensemble to decoding, with each column corresponding to a different ensemble. Data have been normalized to the maximum decoding strength by an ensemble for each variable to allow comparison of ensemble decoding strength across each variable. \**p* < 0.05; \*\**p* < 0.01; \*\*\**p* < 0.001 for post-hoc comparisons.

Specific neurons could be reliably identified across days based on structure and relative position within each FOV (Figure 4A). Thus, we performed single cell tracking across learning and after learning to determine neuronal response evolution and maintenance within all recorded neurons and within each neuronal ensemble. Pearson’s regression analyses were used to compare activity between each of the tracked behavioral sessions (Figure 4C-D, 5B). Cell tracking was performed by a student blinded to experimental conditions to reduce the potential for experimenter-related biases.

**Figure 4.**
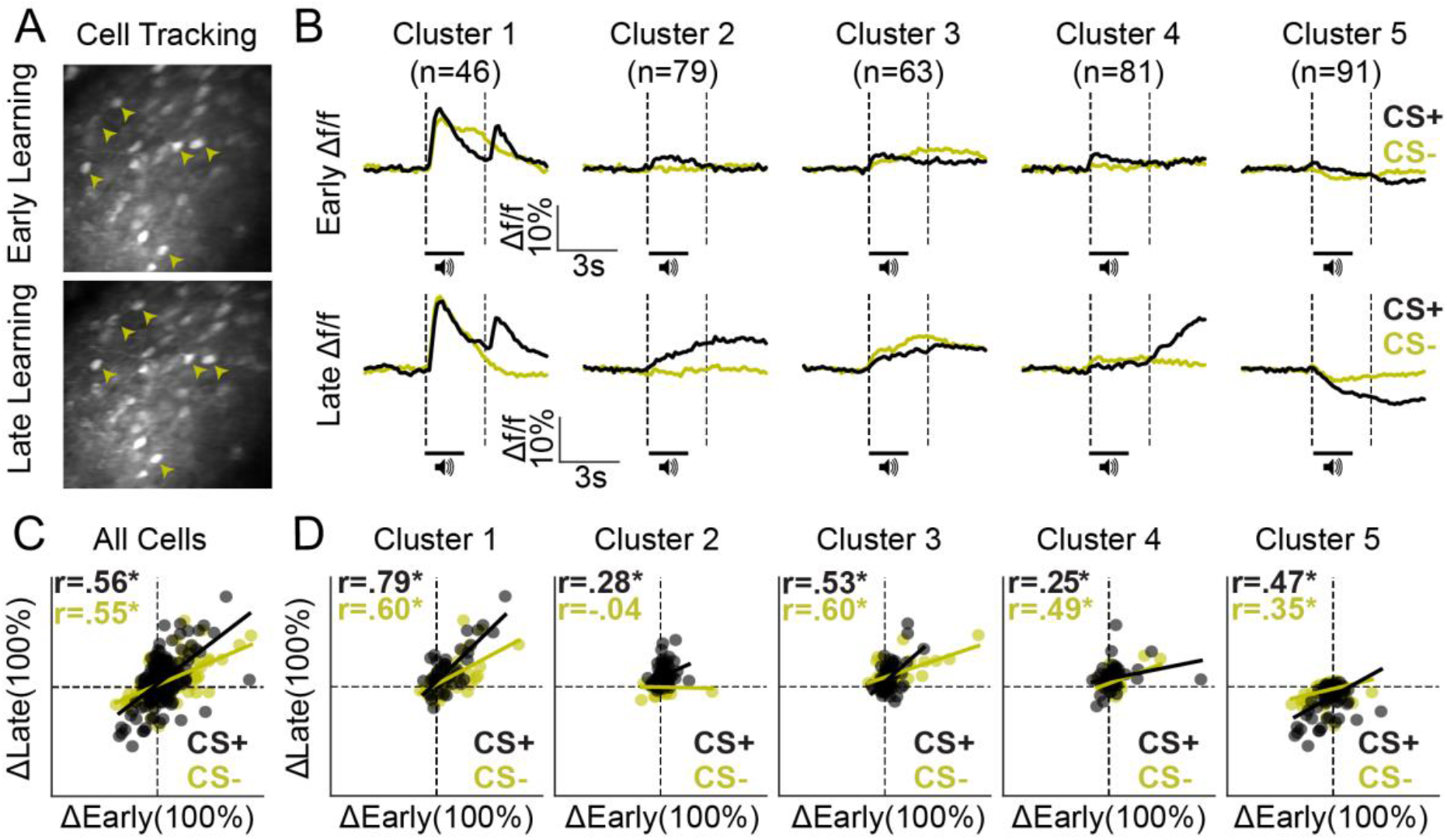
Ensemble-specific activity dynamics differentially evolve across learning. (**A**) Example FOV showing the same neurons (arrowheads) from early (top) and late (bottom) in learning sessions, tracked across days. (**B**) Mean activity traces during CS+ and CS- trials for each ensemble early (top row) and late (bottom row) in learning. (**C**) Correlation plot displaying the change in activity (baseline vs. cue/reward period) of all tracked neurons during CS+ and CS- trials early and late in learning. (**D**) Correlation plots separated by cluster displaying the change in activity (baseline vs. cue/reward period) during each trial early and late in learning. Pearson-R values are displayed in the top left corner for all cells and for each ensemble (C-D). \**p* < 0.05.

**Figure 5.**
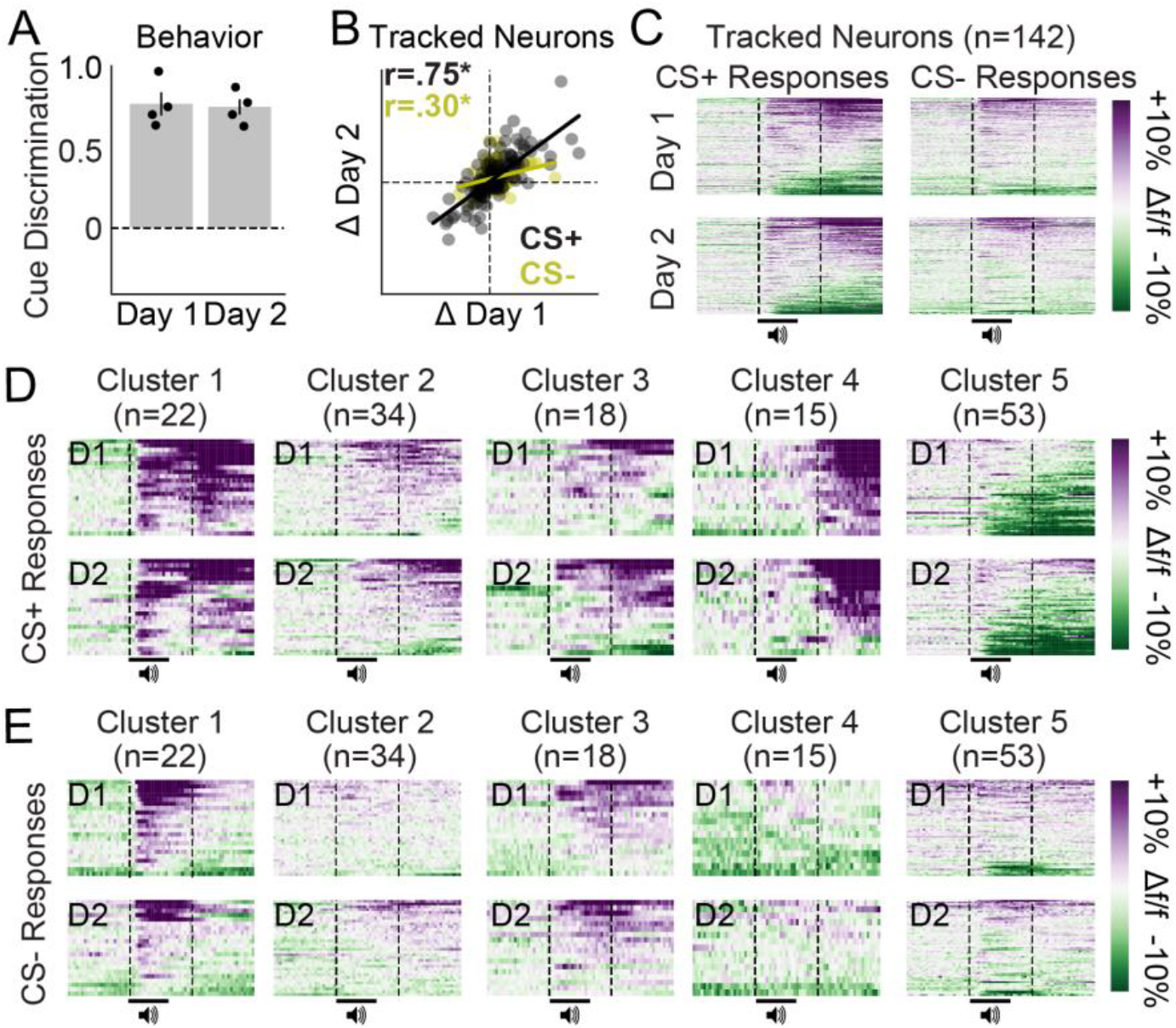
Ensemble-specific activity dynamics are maintained across days after learning. (**A**) Cue discrimination scores (auROC; CS+ vs. CS-) showing similar behavior across days after learning. (**B**) Correlation plots displaying the change in activity (baseline vs. cue/reward period) during CS+ and CS-trials for all tracked neurons during two imaging sessions late in learning. These responses were highly correlated (Pearson-R *p*-values < 0.001). (**C**) Activity heat maps for all tracked neurons separated by cue (columns) and day (rows) reveal similar population dynamics across days after learning. (**D-E**) Activity heat maps for each ensemble separated into CS+ trials (D) and CS-trials (E) confirm similar activity patterns across days after learning.

## RESULTS

### Unique excitatory neuronal ensembles in dmPFC during sucrose seeking

We employed a Pavlovian sucrose conditioning task wherein head-restrained mice were trained to associate one tone conditioned stimulus (CS+), but not another (CS-), with the delivery of a liquid sucrose reward (**Figure 1A-B**). Mice readily acquired this task, showing conditioned licking behavior between the CS+ offset and reward delivery (trace interval), but not the CS- offset and reward omission (**Figure 1C-D**). An independent t-test showed that cue discrimination scores (auROCs) from late in learning behavioral sessions were significantly higher than scores from early in learning behavioral sessions (t_47_ = 23.16; *p* < 0.001; n = 10 mice and 28 FOVs early in learning; n = 7 mice and 21 FOVs late in learning). Furthermore, a two-way ANOVA revealed a significant cue by session interaction for conditioned licking behavior (Δ lick rate; F_1,47_ = 165.1; p < 0.001), and post-hoc tests confirmed that mice licked significantly more during CS+ trials during late in learning sessions as compared with CS-trials during both sessions (*p*-values < 0.001) and CS+ trials during early in learning sessions (*p*-value < 0.001). Thus, conditioned licking behavior for the CS+, but not CS-, developed throughout training revealing that the cue-reward association had been acquired.

We monitored the activity dynamics of putative dmPFC excitatory output neurons throughout the Pavlovian sucrose conditioning task using two-photon calcium imaging (via AAVdj-CaMK2α-GCaMP6s) *in vivo* (**Figure 1E-F**). Similar to previous reports (Otis et al., 2017, 2019), we found that neurons display excitatory and inhibitory responses (n=1511 neurons from 21 FOVs) to each cue and/or to the reward during behavioral sessions after learning (**Figure 1G, 1I**). To better classify these post-learning activity patterns, we used a spectral clustering algorithm (Namboodiri et al., 2019) to isolate unique responses between recorded neurons. The analysis revealed the existence of 5 clusters or “neuronal ensembles” that comprise the majority of response variability (**Figure 1H, 1J**). Each neuronal ensemble displayed a unique activity pattern during the CS+ and/or CS- trials, with qualitative analyses revealing the following dynamics: <u>Cluster 1</u>: excitatory responses during CS+, CS-, and reward delivery (n = 192 neurons from 5/7 mice; 17/21 FOVs), <u>Cluster 2</u>: excitatory responses during CS+ trials (n = 346 neurons from 7/7 mice; 19/21 FOVs), <u>Cluster 3</u>: excitatory responses during CS+ and CS- trials (n = 291 neurons from 7/7 mice; 20/21 FOVs), <u>Cluster 4</u>: excitatory responses during reward delivery (n = 320 neurons from 5/7 mice; 17/21 FOVs), and <u>Cluster 5</u>: inhibitory responses during CS+ trials (n = 362 neurons; 7/7 mice; 21/21 FOVs). Overall, we find the existence of 5 unique ensembles among dmPFC excitatory output neurons, with each of these ensembles displaying unique activity patterns after learning in a Pavlovian sucrose seeking task.

### Select excitatory neuronal ensembles in dmPFC predict behavioral performance during conditioned sucrose seeking

Considering that some neuronal ensembles were absent within particular FOVs during a behavioral session, we determined whether the relative proportion of each neuronal ensemble in a given FOV predicts behavioral performance. To this end, we quantified the proportion of neurons within each ensemble for all behavioral sessions (n = 21 sessions, 21 unique FOVs), and compared those values to cue discrimination licking scores (auROC, CS+ vs. CS-lick rate). Overall, we find that proportion of neurons within Cluster 5 (which showed inhibitory responses during CS+ trials) predicts cue discrimination licking scores (**Figure 2A**). Pearson-R correlation values, found in the inset of each subpanel (Figure 2A), reveal a significant, positive relationship between cue discrimination licking scores and the percentage of neurons in Cluster 5 (the CS+ inhibited ensemble; *p*-value = 0.01). However, there was no significant correlation found for Clusters 1-4 (*p*-values > 0.2). These data are particularly interesting considering previous findings showing that dmPFC PVT neurons, which are also primarily inhibited during CS+ trials, are critical for the acquisition and expression of cue-induced reward seeking (Otis et al., 2017, 2019). These findings suggest that mice with greater numbers of neurons displaying inhibitory CS+ responses after learning, as in Cluster 5, may have improved behavioral performance in the reward seeking task.

We also investigated whether the number of neurons within a particular neuronal ensemble predicted the probability that mice would increase licking during the CS-after learning (auROC, CS- vs. baseline lick rate), which could be considered the initiation of a licking “error”. Overall, we find that the relative proportion of neurons within Cluster 3 (which showed equivalent excitatory CS+ and CS- responses) predicts an increase in licking during CS-trials (**Figure 2B**). Pearson-R correlation values, found in each subpanel (Figure 2B), reveal a significant, positive relationship between the initiation of CS-licking “errors” and the percentage of neurons in Cluster 3 (*p*-value < 0.01). However, there was no significant correlation found for other clusters (*p-*values > 0.23). These data suggest that mice with greater numbers of neurons that display equivalent, excitatory responses to both cues, as in Cluster 3, may be more likely to initiate reward seeking when rewards are not available.

### Excitatory neuronal ensembles in dmPFC display specialized coding during sucrose seeking

We find that dmPFC excitatory neuronal ensembles display unique activity patterns after Pavlovian sucrose conditioning, and that the relative proportion of neurons in each ensemble (specifically, Clusters 3 and 5) can predict behavioral task performance. Despite these findings, whether dmPFC activity patterns can be used to reliably infer environmental or behavioral events during the task is unknown. To this end, we trained a decoder to predict cue, reward, and licking events based on the activity dynamics of all neurons within each FOV (early in learning, n = 28 FOVs, late in learning, n = 21 FOVs). Overall, we find that dmPFC population dynamics within each FOV can be used to detect the presentation of the CS+, CS-, CS+ vs CS- (cue discrimination), reward, and licking rate during late in learning sessions, whereas these activity patterns can be used to predict only the CS+, CS-, and CS+ vs CS- (cue discrimination), but not reward delivery or licking rate during early in learning sessions (**Figure 3A**). ANOVAs, main effects of shuffling: <u>CS+</u>, F_1,92_ = 37.29, *p*-value < 0.001; posthoc *p*-values < 0.05; <u>CS-</u>, F_1,92_ = 22.88, *p*-value < 0.001; posthoc *p*-values < 0.05. <u>CS+ vs CS-</u>, F_1,92_ = 35.94, *p*-value < 0.001; posthoc *p*-values < 0.01; <u>Reward</u>, F_1,92_ = 27.41, *p*-value < 0.001; early in learning posthoc *p*-value > 0.05, late in learning posthoc *p*-value < 0.001; <u>Licking</u>, F_1,92_ = 23.94, *p*-value < 0.001; early in learning posthoc *p*-value > 0.05, late in learning posthoc *p-*value < 0.001. A heatmap illustrating normalized decoding early and late in learning, and the change in that decoding across learning, reveals improved CS+ detection (posthoc *p*-value < 0.05), reward detection (posthoc *p*-value < 0.05), and lick rate prediction across learning (posthoc *p*-value < 0.01; other *p-*values > 0.05; **Figure 3B**). Overall, the activity dynamics of dmPFC excitatory output neurons can be used for cue detection, cue discrimination, reward detection, and prediction of licking after learning. However, activity in these neurons cannot be used to accurately infer reward delivery or licking during sessions early in learning, suggesting that the coding of these variables may be learning dependent.

The population dynamics of dmPFC excitatory output neurons can be used to predict environmental and behavioral factors related to conditioned sucrose seeking, but how unique dmPFC neuronal ensembles contribute to this information coding is unknown. Thus, we next trained a decoder to predict information related to the Pavlovian sucrose seeking task based on the activity dynamics of neurons within each ensemble. Overall, we find superior decoding of the CS+, CS-, CS+ vs CS- (cue discrimination), reward, and conditioned licking in select neuronal ensembles (**Figure 3C, 3D**). <u>CS+</u>: The timing of CS+ presentation could be decoded based on the activity of all cell clusters, although it was best predicted based on the activity of neurons within Cluster 1. <u>CS-</u>: The CS- was significantly predicted by Clusters 1, 3, 4, and 5, and also best predicted based on the activity of Cluster 1. <u>CS+ vs CS-</u>: Activity in Cluster 2, but not other clusters, could be used to significantly discriminate between the CS+ and CS-. <u>Reward</u>: Activity within Clusters 1, 4, and 5 could be used to detect the reward, with activity in Cluster 4 being the best predictor. <u>Licking</u>: Activity in Cluster 5, but not other clusters, could be used to decode conditioned licking. ANOVAs, main effects of shuffling: <u>CS+</u>, F_5,162_ = 14.10, *p*-value < 0.001; posthoc *p*-values < 0.05 for all clusters; <u>CS-</u>, F_5,162_ = 12.72, *p*-value < 0.001; posthoc *p*-values < 0.05 for Clusters 1, 3-5; posthoc *p*-value = 0.84 for Cluster 2. <u>CS+ vs CS-</u>, F_5,162_ = 3.46, *p*-value = 0.005; posthoc *p*-value = 0.004 for Cluster 2, posthoc *p-*values > 0.10 for other clusters; <u>Reward</u>, F_5,162_ = 8.03, *p*-value < 0.001; posthoc *p*-values < 0.007 for Clusters 1, 4, and 5; posthoc p-values > 0.24 for Clusters 2 and 3. <u>Licking</u>, F_5,162_ = 2.91, *p*-value = 0.015; posthoc *p*-value = 0.005 for Cluster 5, posthoc *p*-values > 0.16 for all other clusters. Overall, these data reveal that dmPFC excitatory neuronal ensembles predict select environmental and behavioral factors related to conditioned sucrose-seeking behavior after learning.

### Excitatory neuronal ensembles in dmPFC differentially develop during Pavlovian sucrose conditioning and are stable after learning

Two-photon microscopy enables visual tracking of single, virally labeled neurons across days (Namboodiri et al., 2019; Otis et al., 2017). Thus, we were able to track dmPFC excitatory output neurons to evaluate response evolution across appetitive learning (**Figure 4A**). Overall, we found that neurons in Cluster 1, which show excitatory responses to both cues and to the reward late in learning, also show robust responses to the same stimuli early in learning (**Figure 4B**). In contrast, neurons in Clusters 2-5 did not show obvious responses before learning during CS+ or CS- trials (**Figure 4B**), suggesting that their activity patterns evolved across conditioning and may therefore be reflective of learning. Interestingly, responses during CS+ and CS-trials late in learning were highly predicted by responses early in learning for all cell clusters (**Figure 4C, 4D**). Pearson-R correlation values can be found in the inset of each subpanel (Figure 4C, 4D), and reveal positive correlations during CS+ and/or CS-trials that are significant for each cluster (*denotes *p*-value < 0.05; n = 5 mice, 9 FOVs, 360 tracked neurons). Thus, although responses in Cluster 1 were apparent during both early and late in learning behavioral sessions, responses in all clusters late in learning could be predicted based on early in learning activity dynamics. Whether responses among these neuronal ensembles are maintained after learning, however, remains unclear.

We next tracked the activity dynamics of dmPFC excitatory neurons after learning to determine if the defined neuronal ensembles remain stable or adapt across days (n = 3 mice, 4 FOVs, 142 neurons). Mice showed equivalent behavioral responses during these two late in learning behavioral sessions (**Figure 5A**), as an independent t-test revealed no change in cue discrimination scores across days (t_6_ = 0.22, *p* = 0.84). Neuronal responses during both CS+ and CS-trials were highly correlated across these sessions (**Figure 5B, 5C**; CS+: Pearson-R = 0.75, *p* < 0.001; CS-: Pearson-R = 0.30, *p* < 0.001). Heatmaps for each cluster reveal these highly correlated response patterns during both CS+ trials (**Figure 5D**) and CS-trials (**Figure 5E**). Overall, these data suggest that activity patterns among dmPFC excitatory neuronal ensembles are stable between days after learning.

## DISCUSSION

Here we characterize unique excitatory neuronal ensembles in dmPFC that differentially predict behavioral task performance and encode specialized information related to Pavlovian sucrose conditioning. The responses of each ensemble differentially emerge across learning in a manner that improves the predictive validity of dmPFC population dynamics for deciphering reward delivery, reward-predictive cue presentation, and task-related behavioral output. Considering the stability of dmPFC activity dynamics after learning, our results suggest that each ensemble may be comprised of a unique set of cell types. Future studies that characterize the circuit connectivity, gene expression, and behavioral function of each neuronal ensemble, defined based on *in vivo* activity dynamics, are essential for understanding the dmPFC circuit contributions to reward processing.

Similar to previous studies, we find heterogenous activity patterns among dmPFC excitatory output neurons during reward seeking (Murugan et al., 2017; Otis et al., 2017, 2019; Siciliano et al., 2019), and use a spectral clustering algorithm to isolate five unique neuronal ensembles. Despite these findings, how each neuronal ensemble may be composed of unique cell types – for example, based on projection specificity – remains unclear. Previously, using the same Pavlovian reward-seeking task described here we found that dmPFC NAc neurons are “generally excited” whereas dmPFC PVT are “generally inhibited” following the presentation of a CS+, but not CS-, such that their overall activity patterns fit well within Cluster 2 (dmPFC NAc) and Cluster 5 (dmPFC PVT; Otis et al., 2017). Interestingly, in that study we found that optogenetic inhibition of dmPFC PVT neurons facilitates cue-reward learning, whereas optogenetic activation of the pathway prevents learning and cue-evoked sucrose seeking. These data are consistent with the current findings showing that the proportion of cue-inhibited dmPFC neurons positively predicts behavioral performance in the task (see Figure 2A). Thus, we predict that Cluster 5 is accounted for in part by dmPFC PVT neurons, although further investigation is required to confirm this idea. Considering the heterogeneity found even in projection-specific recording studies in dmPFC (Murugan et al., 2017; Otis et al., 2017, 2019; Siciliano et al., 2019; Vander Weele et al., 2018), it is unlikely that a single projection pathway could be isolated to a single neuronal ensemble. To unravel the circuit connections of these unique cell types, it will therefore be necessary to selectively label neurons based on their *in vivo* activity dynamics, allowing for post-hoc examination of their connectivity. Virally packaged fluorescent proteins that allow light-driven labeling of activated neurons, such as CaMPARI (Fosque et al., 2015), could be useful in this regard but also have limitations that would prevent precision labeling of selected neuronal ensembles (for example, all activated cells would be labeled during UV light delivery, rather than only cells determined to be within a defined ensemble). Thus, development of novel technologies that allow robust labeling of experimenter-selected neurons are critical for identifying the projection profile of unique neuronal ensembles not only in dmPFC, but throughout the brain.

In addition to distinct circuit connectivity patterns, dmPFC neuronal ensembles may display differences in gene expression that could account for their unique activity dynamics. Although little is known about ensemble-specific gene expression in the dmPFC, cortical excitatory projection neurons are thought to express Camk2α (Dittgen et al., 2004), and there is evidence that the immediate early gene NPAS4 is upregulated in reward-responsive, but not aversion-responsive projection neurons (Ye et al., 2016). Additionally, layer-specific gene expression patterns may be present, such as in the case for genes encoding dopamine receptors (Gaspar et al., 1995), nicotinic acetylcholine receptors (Verhoog et al., 2016), noradrenergic receptors (Santana et al., 2013), and more (for review, see Santana and Artigas, 2017). However, recording experiments from subpopulations of dmPFC excitatory output neurons, defined based on gene expression, during reward seeking have been limited. Altogether, a more thorough characterization of ensemble-specific gene expression is needed to ascertain whether genetic differences account for unique activity patterns among dmPFC excitatory neuronal ensembles. Experiments involving single-cell sequencing *ex vivo*, such as patch-*seq* (Cadwell et al., 2016, 2017), could provide gene expression readouts from neuronal ensembles detected *in vivo* to improve our understanding of the gene expression differences that contribute to the evolution of distinct neuronal ensembles.

Our data showing learning-related, stimulus-specific activity patterns among dmPFC excitatory neuronal ensembles is consistent with previous studies. Previous investigations harnessing *in vivo* electrophysiology have found coordinated activity among undefined dmPFC cell populations during reward seeking, for example related to consummatory behavior (licking) in a learning task (Horst and Laubach, 2013), initiation of complex behavioral strategies (Powell and Redish, 2014), recently committed errors in behavioral responding (Powell and Redish, 2014), and behavioral actions based on flexible information (such as rule shifting; Bissonette and Roesch, 2015; Del Arco et al., 2017; Durstewitz et al., 2010; Powell and Redish, 2016; Rodgers and DeWeese, 2014). Using waveform matching across days, one study even demonstrated day-to-day stability in behavioral strategy-related firing patterns (Powell and Redish, 2014), similar to another study showing stable *fos* expression, a marker of activated neurons, in dmPFC neurons across days in an appetitive conditioning task (Brebner et al., 2020). Altogether, these data are consistent with our findings showing day-to-day stability in variable-specific encoding patterns among dmPFC neuronal ensembles.

Here we identify several distinct dmPFC excitatory neuronal ensembles during a Pavlovian sucrose-conditioning task. Despite the apparent simplicity of this task, our findings reveal complex and specialized coding patterns among these heterogeneous neuronal ensembles, which are unlikely to be specific to one particular projection pathway or gene expression profile. Finally, our data suggest that unique aspects of reward seeking may be controlled by distinct neuronal ensembles. Functionally targeting each neuronal ensemble independently, such as through ensemble-specific single cell optogenetic experiments, is therefore critical for understanding how these complex coding patterns control behavioral output (Marshel et al., 2019). Although our results improve our working knowledge of the unique excitatory neuronal ensembles within dmPFC during conditioned reward seeking, they also highlight critical gaps in the field of neuroscience that are important to resolve through new and emerging neurotechnologies.

## Acknowledgements

The authors would like to thank Vijay M.K. Namboodiri and Garret D. Stuber for creating and sharing clustering codes for imaging analysis.

